# MNI-FTD Templates: Unbiased Average Templates of Frontotemporal Dementia Variants

**DOI:** 10.1101/2020.11.25.398305

**Authors:** Mahsa Dadar, Ana L. Manera, Vladimir S. Fonov, Simon Ducharme, D. Louis Collins

## Abstract

Standard anatomical templates are widely used in human neuroimaging processing pipelines to facilitate group level analyses and comparisons across different subjects and populations. The MNI-ICBM152 template is the most commonly used standard template, representing an average of 152 healthy young adult brains. However, in patients with neurodegenerative diseases such as frontotemporal dementia (FTD), the high levels of atrophy lead to significant differences between the brain shape of the individuals and the MNI-ICBM152 template. Such differences might inevitably lead to registration errors or subtle biases in downstream analyses and results. Disease-specific templates are therefore desirable to reflect the anatomical characteristics of the populations of interest and to reduce potential registration errors when processing data from such populations.

Here, we present MNI-FTD136, MNI-bvFTD70, MNI-svFTD36, and MNI-pnfaFTD30, four unbiased average templates of 136 FTD patients, 70 behavioural variant (bv), 36 semantic variant (sv), and 30 progressive nonfluent aphasia (pnfa) variant FTD patients as well as a corresponding age matched average template of 133 healthy controls (MNI-CN133), along with probabilistic tissue maps for each template. The public availability of these templates will facilitate analyses of FTD cohorts and enable comparisons between different studies in a common standardized space appropriate to FTD populations.

## Background and Summary

Most brain image processing pipelines use average templates as a target for registration, to enable use of prior anatomical information and to obtain a common coordinate system based on which they can perform group level analyses and comparisons (Ashburner et al., 2014; Aubert-Broche et al., 2013; Jenkinson et al., 2012; Mateos-Pérez et al., 2018). The MNI-ICBM152 is the most commonly used average template in the neuroimaging literature. However, in certain populations such as pediatric cohorts or patients with neurodegenerative diseases, the variations between the individual brains and the standard MNI-ICBM152 template of young adults might hinder registration accuracy and lead to increase in registration errors (Fonov et al., 2011). In addition, an ill-matched template may give rise to subtle biases in registration that affect processed results. Age appropriate and disease specific templates are therefore desirable not only to reflect the overall anatomical differences between the populations of interest and average young healthy adult brains, but also to reduce potential registration errors and biases when processing data from such populations.

Frontotemporal dementia (FTD) is a clinical categorization describing a heterogenous group of progressive neurodegenerative clinical syndromes associated with atrophy of the frontal and/or anterior temporal lobes. FTD represents about 5% of all cases of dementia in unselected autopsies and, together with Alzheimer’s Disease (AD), it is one of the most common causes of early-onset dementia (Onyike and Diehl-Schmid, 2013). FTD is divided into three major clinical syndromes: the behavioral variant (bvFTD) characterised by prominent early behavioral and personality changes, and the two language variants: the semantic variant (svFTD) and the non-fluent primary progressive aphasia (pnfaFTD). The language variants, also known together as primary progressive aphasias, show language deficits (production, naming, syntax or comprehension) as the main symptom at disease onset without remarkable behavioral disturbance (Bang et al., 2015). A third language variant, the logopenic PPA characterized by prominent hesitations and word retrieval problems, is often included in the FTD umbrella, but it is pathologically most often associated to AD as opposed to frontotemporal lobar degeneration.

Due to the remarkable heterogeneity in FTD neuropathology, as well as the syndromic overlap with other dementias and psychiatric disorders, a confirmed diagnosis within the FTD spectrum is often difficult to achieve in the absence of a dominant genetic mutation (which represents the majority of cases). Therefore, brain imaging with magnetic resonance imaging (MRI) is paramount to increase the level of diagnostic confidence. While it is still not part of standard clinical practice, the potential value of morphometric MRI analysis for diagnostic purposes has been extensively demonstrated (Manera et al., 2019; McCarthy et al., 2018). Hence, improving registration accuracy using disease specific templates could allow better characterization of the pattern of atrophy and its change over time, which could be a valuable resource for both single subject diagnosis and surrogate imaging outcome in clinical trials.

We present MNI-FTD136, MNI-bvFTD70, MNI-svFTD36, and MNI-pnfaFTD30, unbiased average templates of the entire FTD cohort as well as templates of the three variants of FTD, respectively. We also present an average template of age-matched control participants scanned with similar parameters (MNI-CN133) at an isotropic resolution of 1×1×1 mm^3^. We also include their corresponding probabilistic tissue maps for grey matter (GM), white matter (WM), and cerebrospinal fluid (CSF), generated through automatic segmentation. The public availability of these templates will facilitate analysis of FTD cohorts and enable comparisons between different studies in a common standardized space appropriate to FTD populations.

## Methods

### Data

The frontotemporal lobar degeneration neuroimaging initiative (FTLDNI) was funded through the National Institute of Aging and started in 2010. The primary goals of FTLDNI are to identify neuroimaging modalities and methods of analysis for tracking frontotemporal lobar degeneration (FTLD) and to assess the value of imaging versus other biomarkers in diagnostic roles. FTLDNI is the result of collaborative efforts at three sites in North America (site 1, site 2 and site 3). For up-to-date information on participation and protocol, please visit: http://memory.ucsf.edu/research/studies/nifd. Data was accessed and downloaded through the LONI platform in August 2018. We included baseline data from FTD (N_bvFTD_=70, N_svFTD_=36, N_pnfaFTD_=30) patients and age-matched control participants (N_Control_=133) from the FTLDNI database who had T1-weighted (T1w) MRI scans available. All subjects provided informed consent and the protocol was approved by the institution review board at all sites. Table 1 provides the demographic information for the participants in each group.

**Table 1.** Demographic characteristics of the FTLDNI participants. Data are numbers (N) or mean ± standard deviation. CDR=Clinical Dementia Rating.

| Measure | MNI-bvFTD70 | MNI-svFTD36 | MNI-pnfaFTD30 | MNI-CN133 |
| --- | --- | --- | --- | --- |
| N | 70 | 36 | 30 | 133 |
| N<sub>Female</sub> | 26 | 16 | 15 | 56 |
| Age | 61.71 ± 6.25 | 62.82 ± 6.26 | 67.76 ± 7.78 | 63.94 ± 7.56 |
| Education | 15.77 ± 3.31 | 17.26 ± 3.05 | 16.03 ± 2.88 | 17.62 ± 1.86 |
| CDR | 1.15 ± 0.63 | 0.70 ± 0.32 | 0.56 ± 0.59 | 0.02 ± 0.11 |

### Preprocessing

All baseline T1w scans were pre-processed in three steps: image denoising (Coupe et al., 2008), intensity non-uniformity correction (Sled et al., 1998), and image intensity normalization into a 0-100 range. The pre-processed images were then linearly (Dadar et al., 2018) registered to the MNI-ICBM152-2009c template (Manera et al., 2020). Brain extraction was then performed using the registered images (Eskildsen et al., 2012). The quality of the registrations and brain masks were visually assessed to ensure they were accurate.

### Template Generation

A previously validated method was used to generate unbiased average templates for the entire FTD cohort, as well as the bvFTD, svFTD, pnfaFTD subgroups and the age-matched control participants (Fonov et al., 2011, 2009). In short, the method implements a hierarchical nonlinear registration procedure using Automatic Nonlinear Image Matching and Anatomical Labelling (ANIMAL) (Collins and Evans, 1997), reducing the step size at each iteration until convergence is reached. This use of iterative nonlinear registrations in the template generation process leads to average brains that reflect the anatomical characteristics of the population of interest while achieving higher levels of anatomical detail (Fonov et al., 2009). Figure 1 presents axial, sagittal, and coronal slices of each template covering the brain, overlaid by the tissue contours of the MNI-CN133 template to highlight their differences.

**Figure 1.**
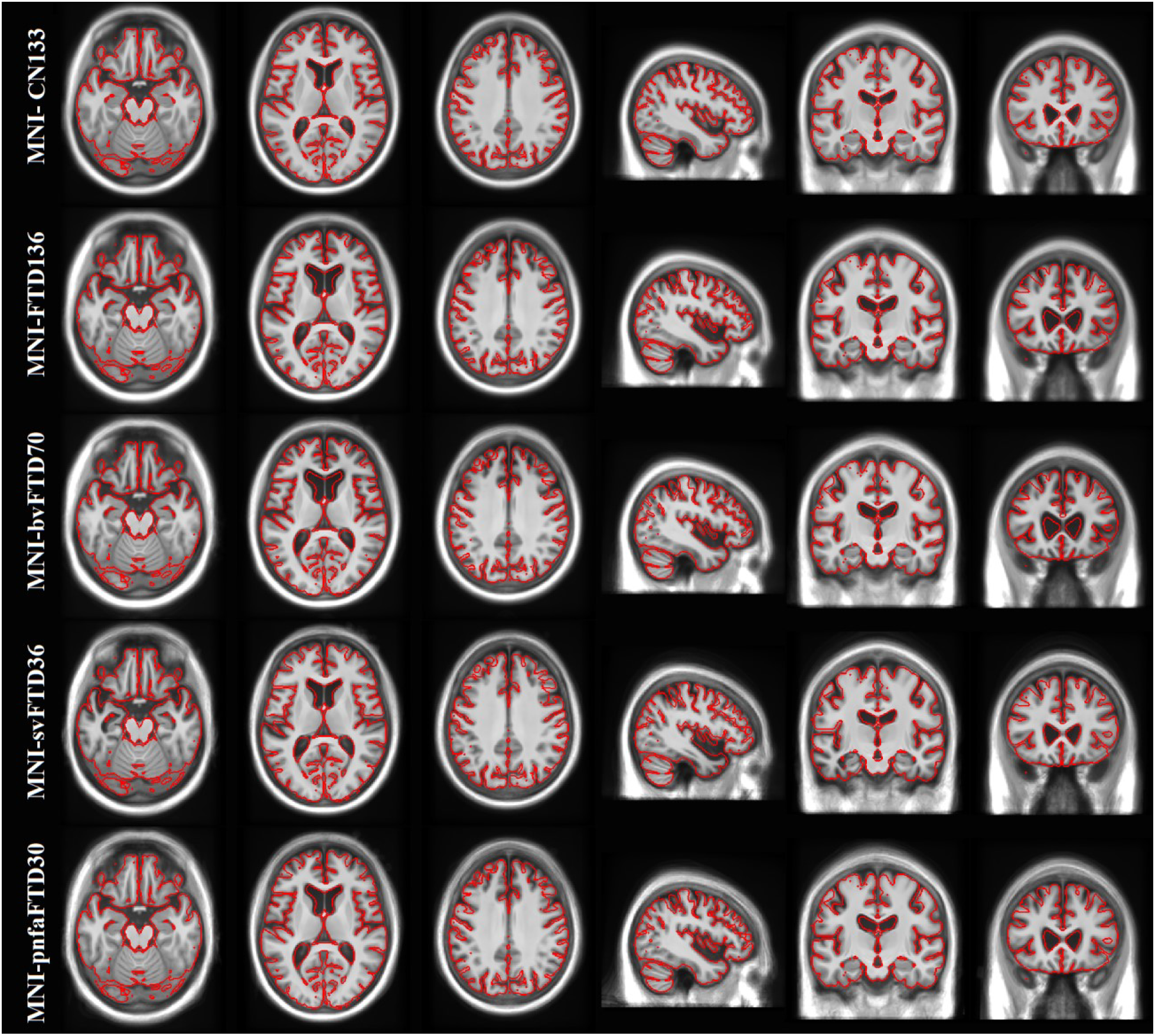
MNI-CN133, MNI-FTD136, MNI-bvFTD70, MNI-svFTD36, and MNI-pnfaFTD30 average templates, overlaid by the contours of the MNI-CN133 template. The figure shows the predominant anterior frontal atrophy compared to controls, which is more evident for the MNI-bvFTD70, whilst MNI-svFTD36 template shows preponderant left anterior temporal atrophy.

The FTD template shows bilateral but asymmetric fronto-insular, anterior temporal and lateral temporal atrophy compared to the age-matched healthy controls (MNI-CN133 template). The bvFTD template shows a similar pattern, but with more evident atrophy on subcortical structures together with greater ventricle enlargement, mainly in the frontal horns of the lateral ventricles. A predominantly left sided temporal atrophy pattern with corresponding ventricular enlargement of the temporal horns is shown on the svFTD template. Finally, the pnfaFTD presents with preponderantly left frontal atrophy with evident enlargement of the frontal horns of the lateral ventricles, though less significant than bvFTD.

### Tissue Maps

Cortical GM, WM, and CSF were automatically segmented for each individual subject using the BISON tissue classification tool, developed and extensively validated for use in multi-center and multi-scanner datasets of aging and neurodegenerative diseases (Dadar and Collins, 2020; Dadar and Duchesne, 2020). Deep GM structures (i.e. putamen, caudate, thalamus, and pallidum) were also segmented using a previously validated deep convolutional neural network (CNN) method (Novosad et al., 2020). The resulting GM, WM, and CSF segmentations from each subject were then nonlinearly resampled to their appropriate templates using the final subject-to-template transform computed in the creation of the unbiased template (e.g. the tissue labels from bvFTD patients were aligned to the MNI-bvFTD70 template). Probabilistic tissue maps were then generated by averaging the nonlinearly registered tissue labels for each cohort. The quality of the segmentations was visually assessed to ensure that only correctly segmented cases were used to create the probabilistic tissue maps. Figures 2-4 show the tissue maps overlaid on their corresponding templates for GM, WM, and CSF.

**Figure 2.**
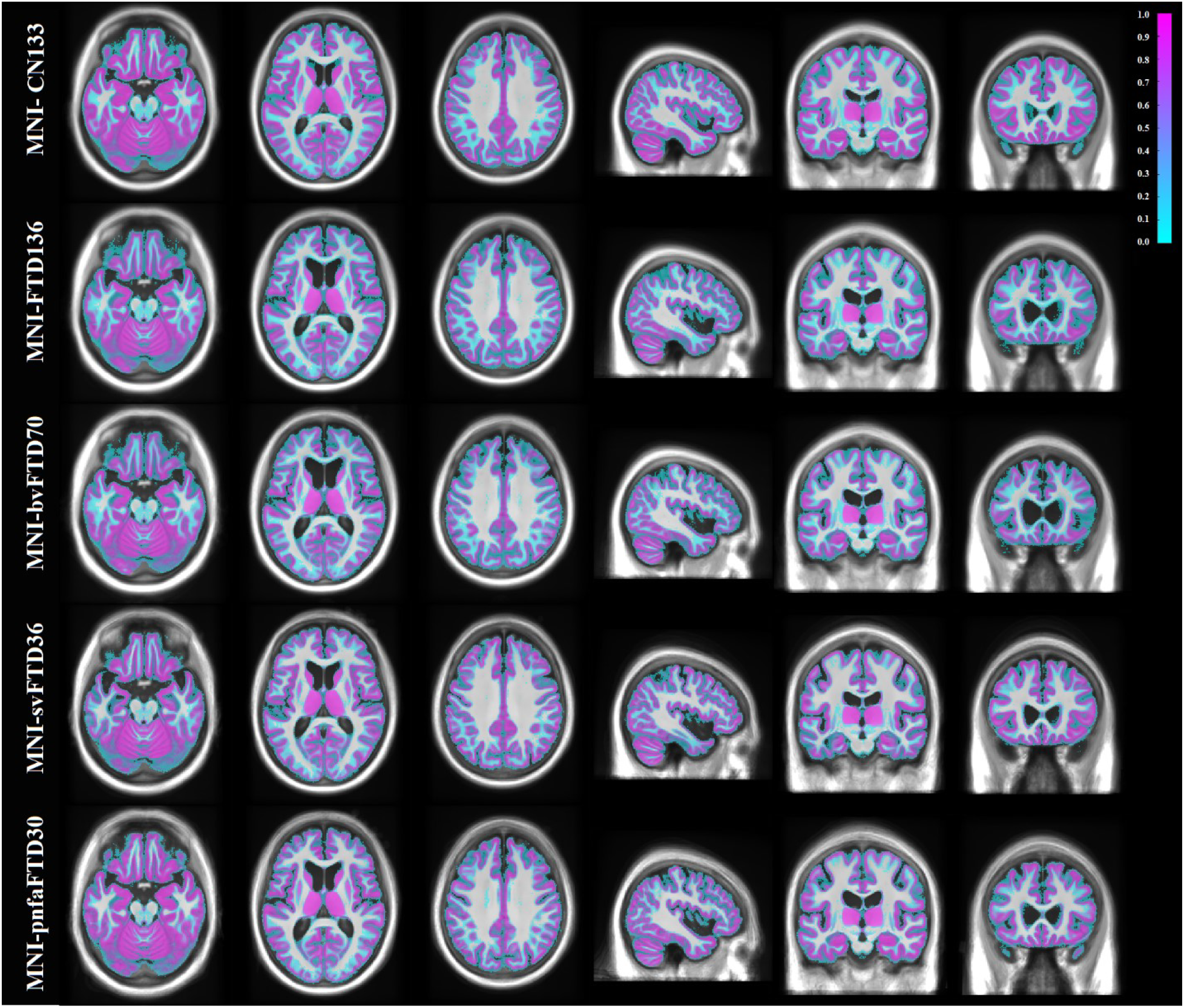
Grey matter probability maps overlaid on the average templates.

**Figure 3.**
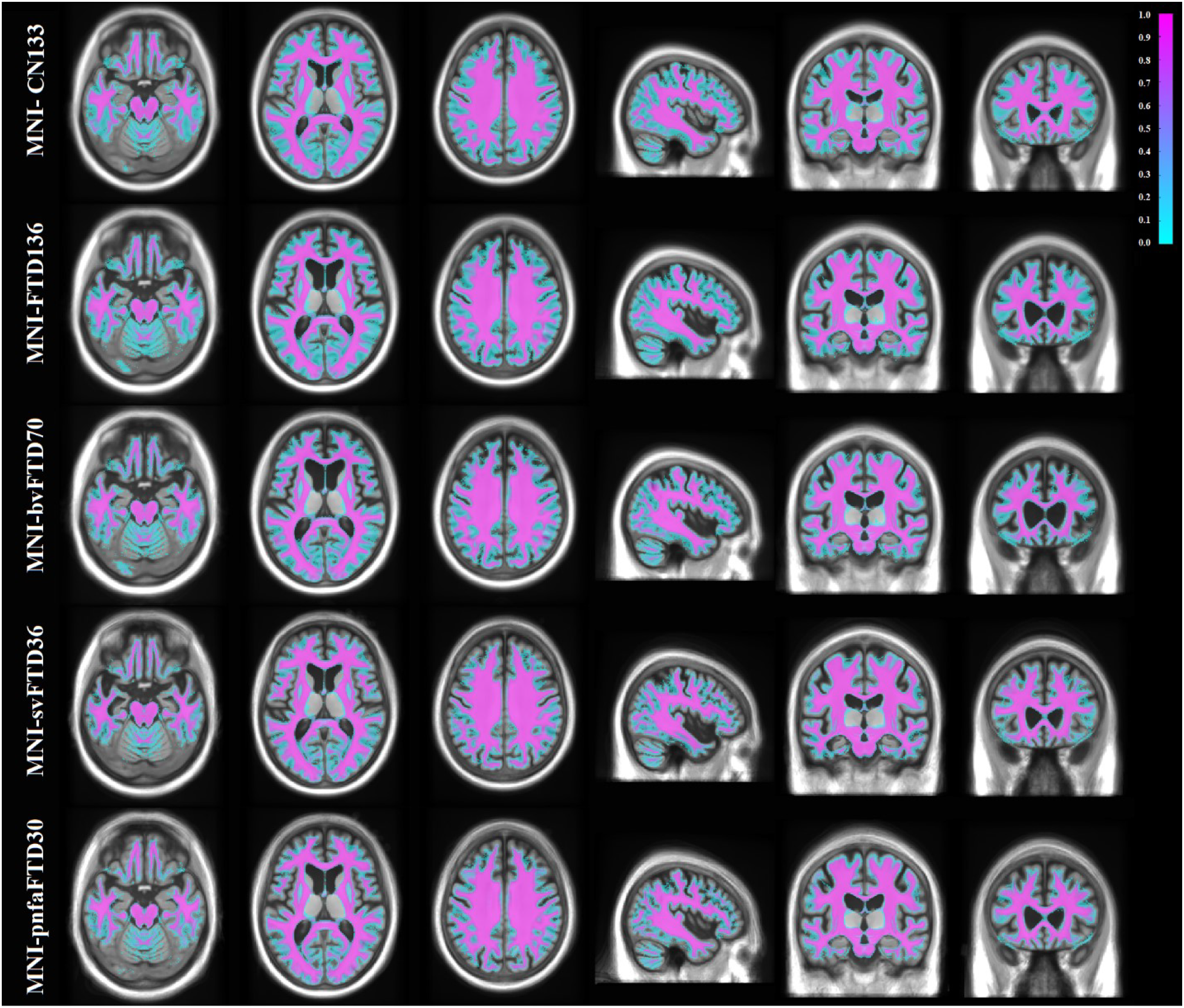
White matter probability maps overlaid on the average templates.

**Figure 4.**
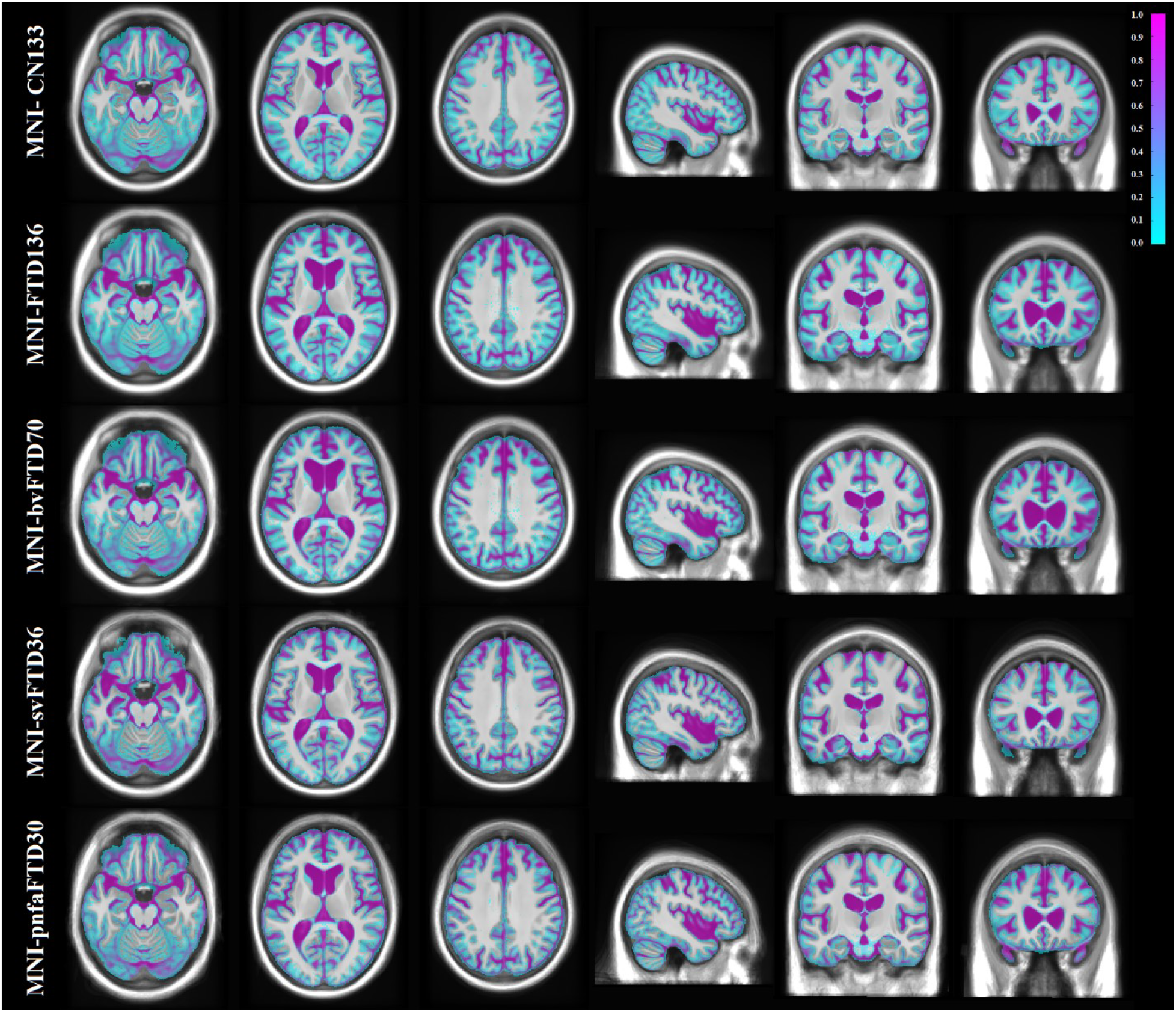
CSF probability maps overlaid on the average templates.

### Data Records

The five average templates (i.e. MNI-FTD136, MNI-bvFTD70, MNI-svFTD36, MNI-pnfaFTD30, and MNI-CN133) as well as their corresponding tissue maps are available at G-Node (https://gin.g-node.org/anamanera/MNI-FTD_Templates.git) and http://nist.mni.mcgill.ca/?p=904. All data are available in compressed MINC (Neelin et al., 1998; Vincent et al., 2016) and NIfTI formats.

### Technical Validation

Prior to generating the average templates, the quality of the images (e.g. presence of image artifacts such as motion) as well as the linear and nonlinear registrations was visually assessed and 16 cases (2 CN, 7 bv FTD, 3 svFTD, and 4 pnfaFTD) that did not pass this quality control step were discarded. Similarly, 10 cases (4 CN, 4 bv FTD, 1 svFTD, and 1 pnfaFTD) failed quality control for tissue segmentation step and were discarded before generating the tissue probability maps.

To further demonstrate the structural differences between the templates, the FALCON cortical surface extraction tool was applied to each template, and the relative cortical thickness difference between each of the FTD templates and MNI-CN133 was calculated (Vladimir S. FONOV, 2020). Similarly, to demonstrate the differences between the templates in the deep gray matter and white matter areas, deformation based morphometry maps were generated based on nonlinear registrations between each FTD template and the MNI-CN133 template (Ashburner et al., 2000). Figure 5 shows the percentage of difference in cortical thickness between MNI-bvFTD70, MNI-svFTD36, and MNI-pnfaFTD30 versus MNI-CN133 cortical surfaces. Colder colors indicate thinner cortex in comparison with MNI-CN133 template. Similarly, Figure 6 shows the regional differences between MNI-bvFTD70, MNI-svFTD36, and MNI-pnfaFTD30 versus MNI-CN133 across the entire brain. Colder colors indicate smaller areas (i.e. shrinkage or more atrophy), and warmer colors (e.g. in the ventricular and sulci regions) indicate larger areas (i.e. expansion) in comparison with MNI-CN133 template.

**Figure 5.**
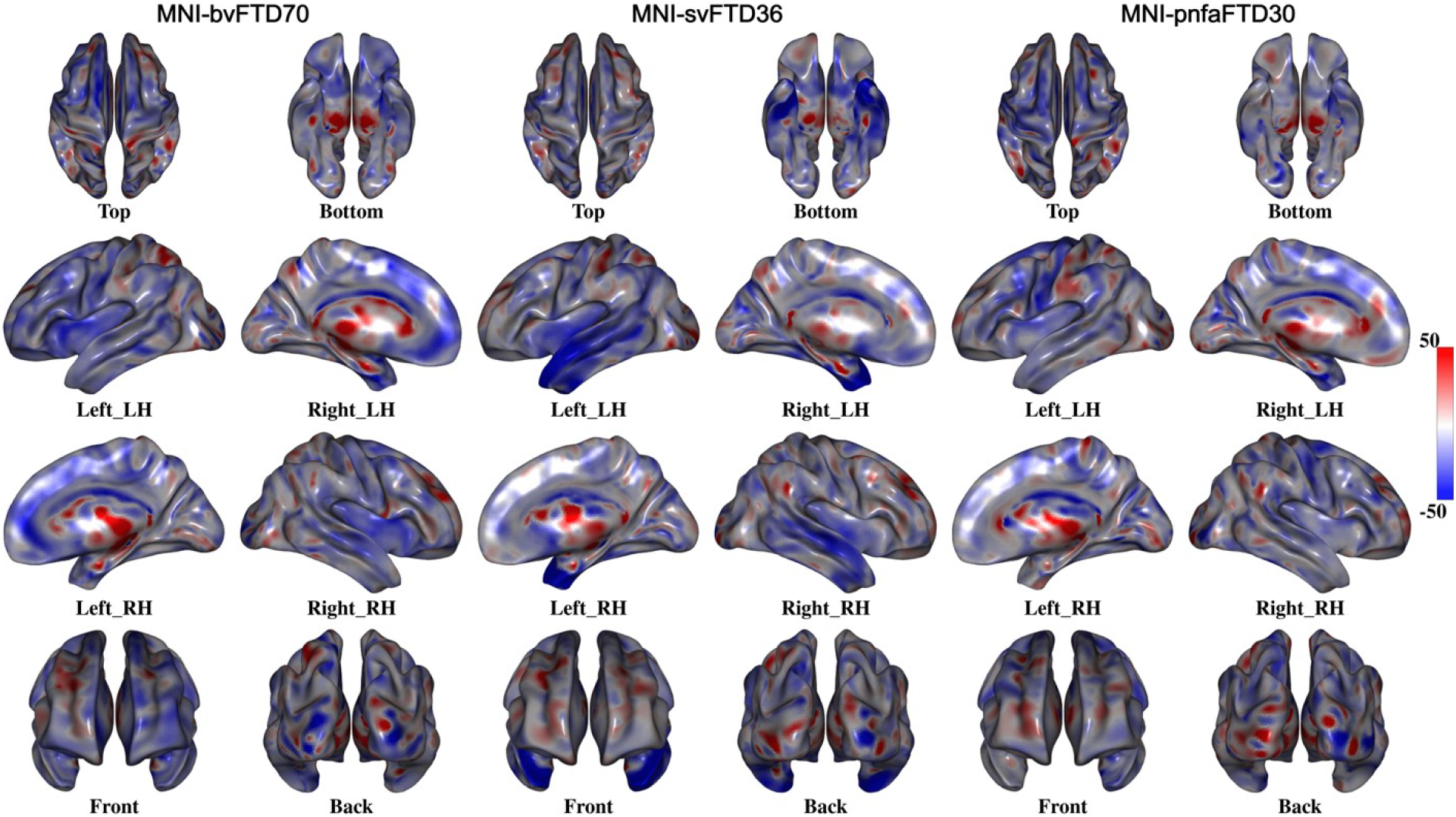
Percentage of cortical thickness difference between MNI-bvFTD70, MNI-svFTD36, and MNI-pnfaFTD30 versus MNI-CN133 templates. MNI-bvFTD70 template shows the largest amount of frontal lobe cortical atrophy. MNI-svFTD36 shows the greatest amount of bi-lateral temporal lobe cortical atrophy compared to the two other FTD variants. MNI-pnfaFTD30 shows left sided cortical thinning in prefrontal and Broca’s areas.

**Figure 6.**
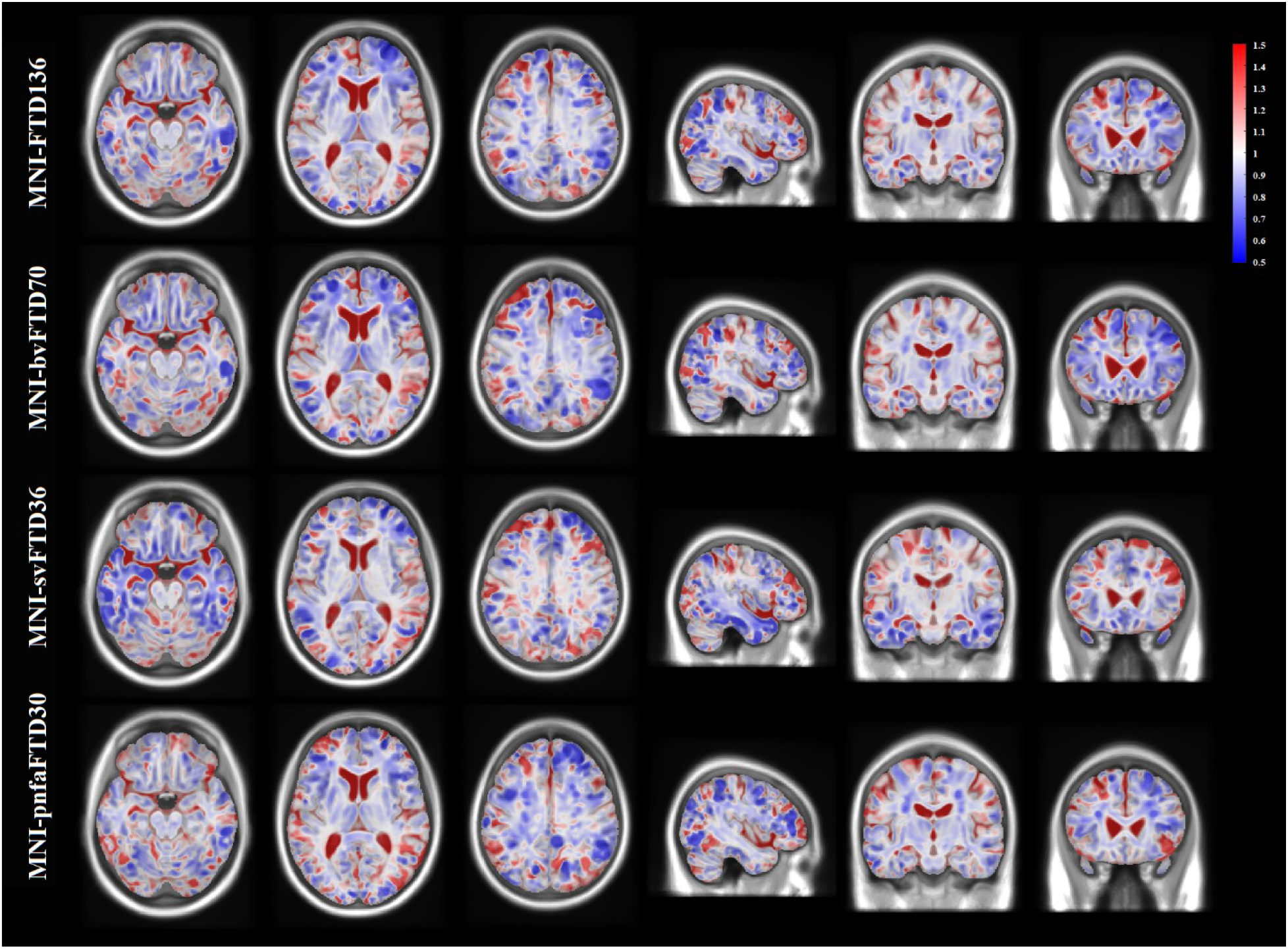
Deformation based morphometry difference between MNI-bvFTD70, MNI-svFTD36, and MNI-pnfaFTD30 versus MNI-CN133 templates. All FTD templates show increased ventricular and sulcal spaces. MNI-bvFTD70 shows predominant bilateral anterior frontal shrinkage. MNI-svFTD36 shows asymmetric in lateral and anterior temporal lobe atrophy. MNI-pnfaFTD30 shows asymmetric frontal lobe atrophy.

Comparing the MNI-bvFTD70 group against the controls, the greatest difference in cortical thickness was located in frontal lobes, bilaterally as well as the temporal poles (specially in medial frontal and dorsolateral prefrontal areas). In MNI-svFTD36, however, cortical thinning is limited to the anterior and lateral temporal lobe, predominantly on the left side. Finally, MNI-pnfaFTD30 shows left sided cortical thinning in prefrontal and Broca’s areas.

Correspondingly, deformation-based morphometry maps show an overall pattern of atrophy in MNI-FTD136, more evident in right anterior frontal and lateral temporal regions. While MNI-bvFTD70 demonstrate a predominant bilateral anterior frontal shrinkage, MNI-pnfaFTD30 shows asymmetric frontal atrophy. As expected, in MNI-svFTD36, smaller areas were located asymmetrically in lateral and anterior temporal lobes. In addition, ventricular and sulci enlargement is also shown for all the templates.

## Abbreviations

bvFTD: Behavioural variant frontotemporal dementia
CSF: Cerebrospinal fluid
FTD: Frontotemporal dementia
GM: gray matter
MRI: magnetic resonance imaging
MNI: Montreal Neurological Institute
pnfaFTD: progressive nonfluent aphasia frontotemporal dementia
svFTD: Semantic variant frontotemporal dementia
T1w: T1-weighted
WM: white matter

## Code Availability

The scripts for generating unbiased average templates, tissue classification, and FALCON are publicly available at https://github.com/vfonov/nist_mni_pipelines, http://nist.mni.mcgill.ca/?p=2148, and https://github.com/philnovv/CNN_NeuroSeg/, and https://github.com/NIST-MNI/falcon.

## Authors Contributions

**Mahsa Dadar**: Study concept and design, analysis of the data, drafting and revision of the manuscript.

**Ana L. Manera**: Study concept and design, interpretation of the data, drafting and revision of the manuscript.

**Vladimir Fonov**: Study concept and design, interpretation of the data, revising the manuscript.

**Simon Ducharme**: Study concept and design, interpretation of the data, revising the manuscript.

**D. Louis Collins**: Study concept and design, interpretation of the data, revising the manuscript.

## Acknowledgements

MD is supported by a scholarship from the Canadian Consortium on Neurodegeneration in Aging (CCNA) as well as an Alzheimer Society Research Program (ASRP) postdoctoral award. AM and SD are also supported by the FTD team from CCNA. The Consortium is supported by a grant from the Canadian Institutes of Health Research with funding from several partners including the Alzheimer Society of Canada, Sanofi, and Women’s Brain Health Initiative.

Data collection and sharing for this project was funded by the Frontotemporal Lobar Degeneration Neuroimaging Initiative (National Institutes of Health Grant R01 AG032306). The study is coordinated through the University of California, San Francisco, Memory and Aging Center. FTLDNI data are disseminated by the Laboratory for Neuro Imaging at the University of Southern California.

